# Quantifying the oxygen preferences of bacterial communities using a metagenome-based approach

**DOI:** 10.64898/2026.01.22.701213

**Authors:** Clifton P. Bueno de Mesquita, Elías Stallard-Olivera, Noah Fierer

## Abstract

Oxygen is a primary driver of the distribution and activity of microbial life. Since oxygen levels are often difficult to measure *in situ*, one potential solution is to use bacteria as bioindicators of oxygen levels. As bacteria range from obligate aerobes to obligate anaerobes, quantification of bacterial community oxygen preferences could be used to infer variation in environmental oxygen levels and bacterial metabolic strategies. After using ensemble machine learning to select the 20 most important genes that predict oxygen tolerances in individual bacteria, we established a relationship between the abundance ratio of aerobic: anaerobic indicator genes and the proportional abundance of aerobic bacteria using simulated metagenomes with varying ratios of known aerobic and anaerobic bacteria. We developed a tool, OxyMetaG, that takes metagenomic reads as input, extracts bacterial reads, maps reads to the 20 genes, and predicts the proportion of aerobic versus anaerobic bacteria in any given sample. We tested OxyMetaG on a suite of metagenomes with measured or inferred oxygen levels across a variety of environmental and host-associated samples. To demonstrate the utility of our approach, we applied OxyMetaG to 540 surface soils, showing that surface soils are typically dominated by aerobes, but wetter sites with finer textures have relatively more anaerobes. Lastly, we applied OxyMetaG to 73 human gut samples, showing that in the first three years of life, human guts progress from having up to 61% aerobes to being completely dominated by anaerobes. We expect OxyMetaG to have broad utility for characterizing both modern and ancient environments.

**Importance:** Oxygen is one of the most important environmental variables affecting microbial activity and composition but is often difficult to measure *in situ*. We developed a tool, OxyMetaG, that leverages differences in bacterial gene content across known aerobic and anaerobic taxa to predict the proportion of aerobes and anaerobes in a given sample directly from shotgun metagenomic reads. OxyMetaG works on samples with low sequencing depth and avoids computationally expensive genome assembly, which often captures only a fraction of the microbial community in a given environment. With OxyMetaG, bacteria can be used as bioindicators of oxygen availability over broader time scales than just a single measurement and provide crucial environmental context in cases where oxygen has not or cannot be measured. OxyMetaG is publicly available and can be used to answer a wide variety of ecological questions in both environmental and host-associated systems.

## Introduction

Throughout geologic history, atmospheric oxygen levels have been variable. For the first ∼1.5 billion years that life likely existed on Earth, atmospheric oxygen was limited and anaerobic bacteria predominated (1). Only following the Great Oxidation Event (2) did the atmosphere become oxygenated, with levels over the past ∼2.4 billion years ranging from 15% to 35% before reaching the current concentration of 21% (3). Even across modern environments, oxygen concentrations can be highly variable (e.g., waterlogged hypoxic wetland sediments to oxic surface ocean water). This spatial and temporal variation in oxygen concentrations is often a primary determinant of bacterial distributions and their metabolic activities. Bacteria can differ widely with respect to their oxygen tolerances and preferences with the oxygen levels in any given environment dictating what taxa can thrive in that environment and their potential metabolic strategies. We sought to take advantage of these patterns to develop a predictive tool that can work directly on metagenomic reads (avoiding computationally expensive assembly, binning, and annotation) to predict the percent relative abundances of aerobic and anaerobic bacteria, which in turn can be used to help infer the relative amount of oxygen in any given environmental or host-associated sample.

Bacterial oxygen preferences range from taxa that are obligate anaerobes that are unable to grow in the presence of oxygen and instead generate energy via fermentation or anaerobic respiration, to obligate aerobes that require atmospheric levels of oxygen to sustain growth. Between these end members exist facultative aerobes/anaerobes that can tolerate oxygen or a lack of oxygen and switch their metabolisms accordingly, and microaerophiles, which are defined as organisms that grow at low (below atmospheric, usually 2-10%) oxygen levels (4). In short, bacterial oxygen preferences span a spectrum and, for most cultivated bacteria, we know from experimental assays where they fall along this spectrum. Even for the majority of bacteria that are resistant to cultivation and have not been studied *in vitro*, oxygen preferences can often be inferred from genome-based models, including models that rely on specific sets of genes (5–14) or other genomic attributes such as amino acid frequencies (15, 16). While such genome-based models enable inferences of oxygen preferences for individual taxa, inferring community-level oxygen preferences would require having high-quality genomes for the majority of taxa in a given community, a challenging task in many systems with high levels of bacterial diversity (17). We thus sought to develop a metagenome-based tool that can work directly on metagenomic reads to predict community-level oxygen preferences, namely the percent of aerobic versus anaerobic bacteria in any given sample. Not only would such a metagenome-based model make it possible to quantify oxygen preferences as a community aggregated trait (18), we could also use inferred community-level oxygen preferences as a means to quantify oxygen availability in individual samples, assuming that the oxygen preferences of bacterial communities should, over time, reflect oxygen availability. In this way, microbes would be used as bioindicators of oxygen availability, as has been done previously for other important but hard-to-measure variables such as phosphorus availability (19), heavy metal contamination, and fecal contamination (20).

Despite its importance, oxygen is notoriously hard to measure in environmental and host-associated samples due to a combination of methodological constraints and the fact that oxygen concentrations are often highly variable across space and time. Take, for example, soils, where oxygen concentrations can vary at the millimeter scale, for example between oxic pore spaces and neighboring anoxic microsites in aggregates (21). Soil oxygen concentrations are also constantly in flux as oxygen is consumed by microbial respiration and supplied by diffusion from the atmosphere. Yet understanding soil oxygen is important for understanding microbial ecology, soil health, and soil biogeochemical process rates (22, 23). As another example, measuring oxygen concentrations in vertebrate guts is notoriously challenging as sensors must be placed inside the individual (24). Yet understanding both human and non-human animal gut oxygen concentrations is crucial for understanding digestion, metabolism, and disease (25–27). Thus, it would be beneficial to infer relative oxygen levels in soils, vertebrate guts, and other environmental or host-associated samples directly from the ever-expanding metagenomic datasets now available.

Here, we describe the development and validation of the new tool OxyMetaG (Figure 1). Our goal was to identify a narrow set of genes with high predictive power at the genome level and then use a read-mapping and abundance-ratio methodology to rapidly profile the raw reads of metagenomes, thereby avoiding the limitations of having to assemble high-quality genomes from metagenomes. After comprehensive validation using simulated metagenomes, we tested OxyMetaG on a wide variety of environmental and host-associated sample types that capture broad gradients in oxygen availability, including a subset of samples with measured oxygen concentrations. We then present two example case studies of how OxyMetaG can be used to answer ecological questions in one environmental system (surface soils) and one host-associated system (human guts). For surface soils, demonstrate how OxyMetaG can be used to quantify continental-scale variation in the predicted relative abundances of aerobic bacteria and the climate or edaphic variables associated with this variation. For human guts, we used metagenomic data from selected individuals sampled over time to quantify how the predicted relative abundance of aerobic bacteria changes during the first three years of life. Together these examples demonstrate how OxyMetaG can be used to identify community-level shifts in bacterial oxygen preferences directly from metagenomic data and infer differences in oxygen availability within and across environments.

**Figure 1.**
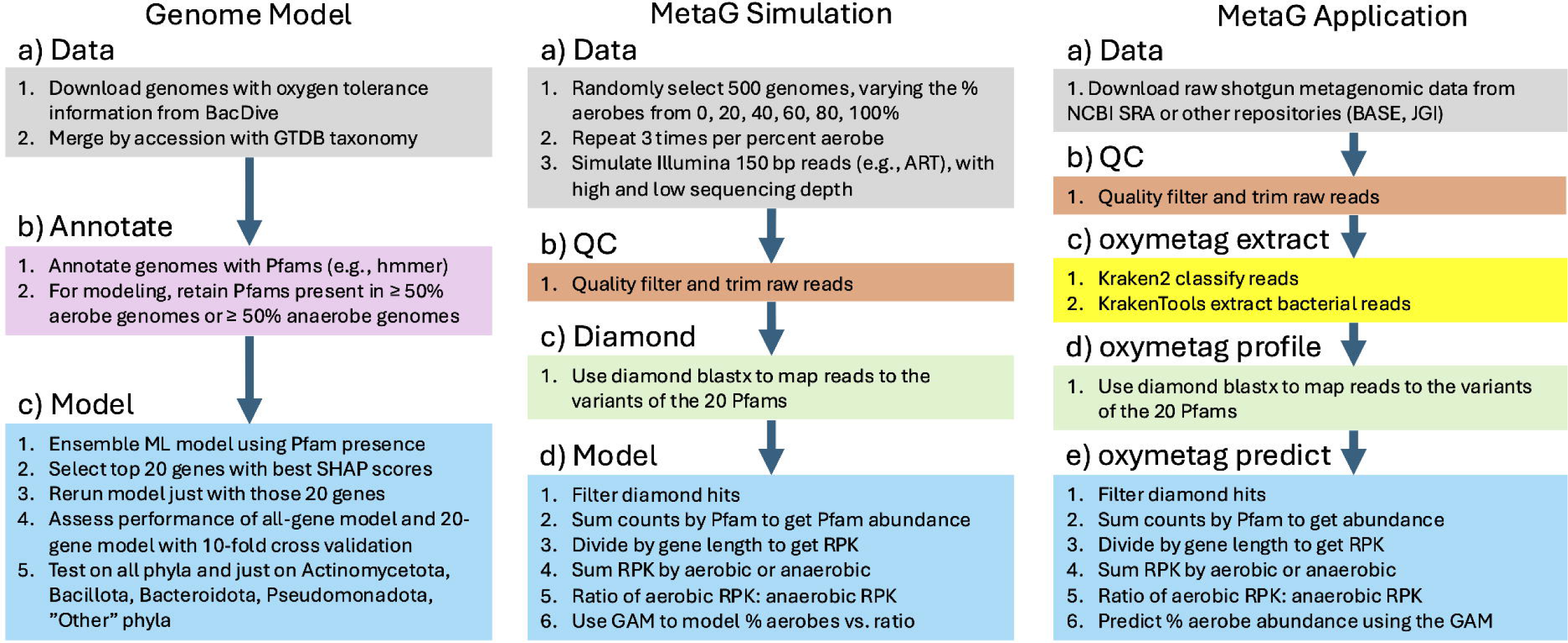
Overview of the development and use of OxyMetaG. 20 predictive Pfams were first selected using a genome-level ensemble machine learning model (Random Forest, CatBoost, XGboost) that utilized oxygen tolerance information from BacDive. Then a relationship between the ratio of selected aerobic indicator genes (n = 13) and anaerobic indicator genes (n = 7) was built based on simulated metagenomes with varying levels of known aerobes and anaerobes. Lastly, the model was tested on 203 metagenomes from various habitats with known or inferred differences in oxygen levels, as well as on 331 surface soil metagenomes from Australia and 209 surface soil metagenomes from the United States. The steps of extracting bacterial reads, running DIAMOND BLASTX, and predicting the percent aerobe relative abundance are implemented in the standalone Python package OxyMetaG, with the functions oxymetag extract, oxymetag profile, and oxymetag predict.

## Results

We identified a set of 20 genes that were highly predictive of oxygen tolerance, including 13 genes associated with aerobes and 7 genes associated with anaerobes (Table 1, Figure S1). The classification model of oxygen tolerance using just those 20 genes had high precision, recall, and F1-score (Table 2), and this was true when tested on all taxa and, the four main phyla (Pseudomonadota, Actinomycetota, Bacillota, Bacteroidota) combined, on each of the main four phyla individually, and on all other rare phyla combined (Table S1). Accuracy per phylum was > 85% for all 31 phyla, and > 93% for 29 of 31 phyla (Table S2).

**Table 1.** The 20 Pfams selected for predicting the oxygen tolerance of individual isolates in BacDive, as well as the Pfam name, whether it is indicative of aerobic or anaerobic metabolism, the percent prevalence in 4227 aerobic bacteria, the percent prevalence in 1293 anaerobic bacteria, and the aggregated SHAP feature importance score.

| **Pfam** | **Name** | **Oxygen** | **Aerobe**<br>**prev.** | **Anaerobe**<br>**prev.** | **FI** |
| --- | --- | --- | --- | --- | --- |
| PF00510 | Cytochrome c oxidase subunit III | aerobic | 94.58 | 12.22 | 0.155 |
| PF00296 | Luciferase-like monooxygenase | aerobic | 89.21 | 13.92 | 0.060 |
| PF16870 | 2-oxoglutarate dehydrogenase C-terminal | aerobic | 96.59 | 17.63 | 0.038 |
| PF00116 | Cytochrome C oxidase subunit II, periplasmic domain | aerobic | 94.42 | 14.23 | 0.038 |
| PF01521 | Iron-sulphur cluster biosynthesis | aerobic | 98.77 | 31.25 | 0.028 |
| PF05425 | Copper resistance protein D | aerobic | 77.88 | 9.59 | 0.023 |
| PF17773 | UPF0176 acylphosphatase like domain | aerobic | 66.57 | 6.19 | 0.022 |
| PF00115 | Cytochrome C and Quinol oxidase polypeptide I | aerobic | 98.11 | 23.28 | 0.020 |
| PF00916 | Sulfate permease family | aerobic | 90.51 | 47.10 | 0.019 |
| PF08530 | X-Pro dipeptidyl-peptidase C-terminal non-catalytic domain | aerobic | 54.67 | 16.71 | 0.019 |
| PF01152 | Bacterial-like globin | aerobic | 74.50 | 6.11 | 0.018 |
| PF05721 | Phytanoyl-CoA dioxygenase (PhyH) | aerobic | 60.70 | 2.24 | 0.017 |
| PF00042 | Globin | aerobic | 53.58 | 5.49 | 0.016 |
| PF01871 | AMMECR1 | anaerobic | 3.38 | 50.66 | 0.074 |
| PF10371 | EKR | anaerobic | 6.60 | 65.82 | 0.058 |
| PF17910 | FeoB cytosolic helical domain | anaerobic | 10.34 | 74.25 | 0.043 |
| PF02579 | Dinitrogenase iron-molybdenum cofactor | anaerobic | 7.55 | 64.89 | 0.041 |
| PF13597 | Anaerobic ribonucleoside-triphosphate reductase | anaerobic | 16.87 | 88.86 | 0.032 |
| PF03063 | Prismane/CO dehydrogenase family | anaerobic | 6.53 | 68.99 | 0.028 |
| PF02906 | Iron only hydrogenase large subunit, C-terminal domain | anaerobic | 1.14 | 58.62 | 0.018 |

**Table 2.** Ensemble model statistics on the 20% holdout test dataset.

|  | **Precision** | **Recall** | **F1-score** | **Support** |
| --- | --- | --- | --- | --- |
| Aerobe | 0.97 | 0.98 | 0.97 | 845 |
| Anaerobe | 0.92 | 0.90 | 0.91 | 259 |
| accuracy |  |  | 0.96 | 1104 |
| macro avg | 0.94 | 0.94 | 0.94 | 1104 |
| weighted avg | 0.96 | 0.96 | 0.96 | 1104 |

In simulated metagenomes with known amounts of aerobic and anaerobic bacterial genomes, the ratio of the summed RPK abundance of the 13 aerobic indicator genes to the 7 anaerobic indicator genes was well correlated with the percent of aerobic bacterial taxa in simulated metagenomes, regardless of sequencing depth (Figure 2). The relationship was non-linear and a generalized additive model was fit to the data (*F* = 1605, R^2^ = 0.99, p < 0.001) that was then used to make predictions on real metagenomes with unknown proportions of aerobic and anaerobic bacteria.

**Figure 2.**
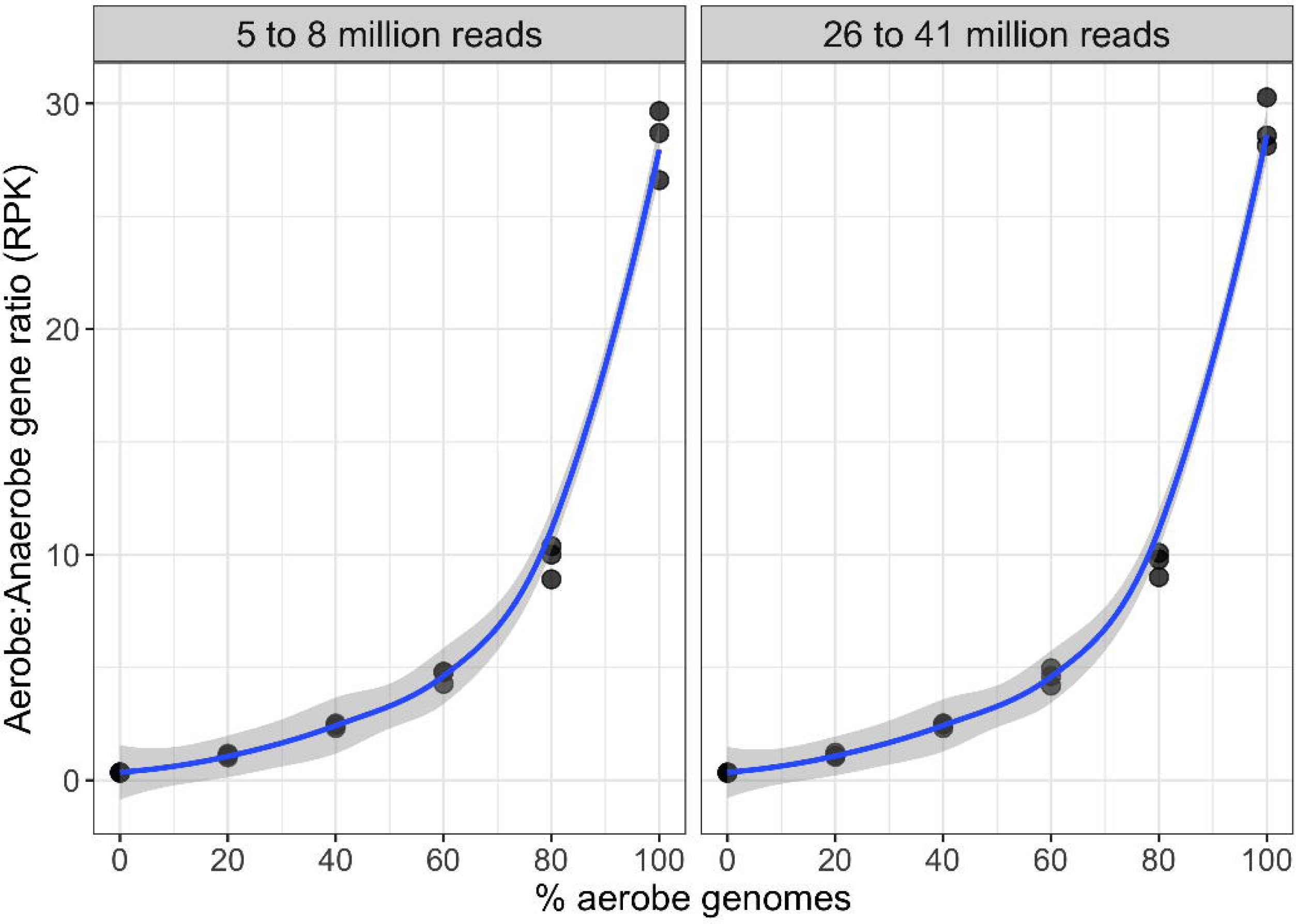
Relationship between the percent of aerobe genomes in simulated metagenomes, and the ratio of the RPK of aerobic indicator genes to anaerobic indicator genes, across two ranges of sequencing depths. The model used for prediction in OxyMetaG is the one trained with data from 26 to 41 million reads.

The predicted percentages of aerobic bacteria closely matched the expected differences in oxygen availability across different habitat types. Samples from cattle rumen, human gut, and marine sediments had the greatest proportion of anaerobes, while surface ocean, grassland soil, and forest soil had significantly more aerobes (Tukey posthoc, p < 0.05, Figure 3). Irrigated temperate cropland soils and tropical forest soils had the greatest relative abundance of anaerobes in any of the upland soil habitats, potentially reflecting greater soil moisture content and consequently lower oxygen concentrations in those soils. Furthermore, in the three datasets where metagenomic data were available with corresponding measurements of oxygen concentrations (Baltic Sea sediment, Black Sea water, Lake Tanganyika water), the predicted relative abundances of aerobic bacteria increased with measured oxygen concentrations (Figure 4). Results were similar when comparing the default cutoffs (percent identity ≥ 60, e-value < 0.001, and bitscore ≥ 50) and more stringent cutoffs (percent identity ≥ 60, e-value < 1e-10, bitscore ≥ 60, and amino acid alignment length ≥ 40) (Figure S2).

**Figure 3.**
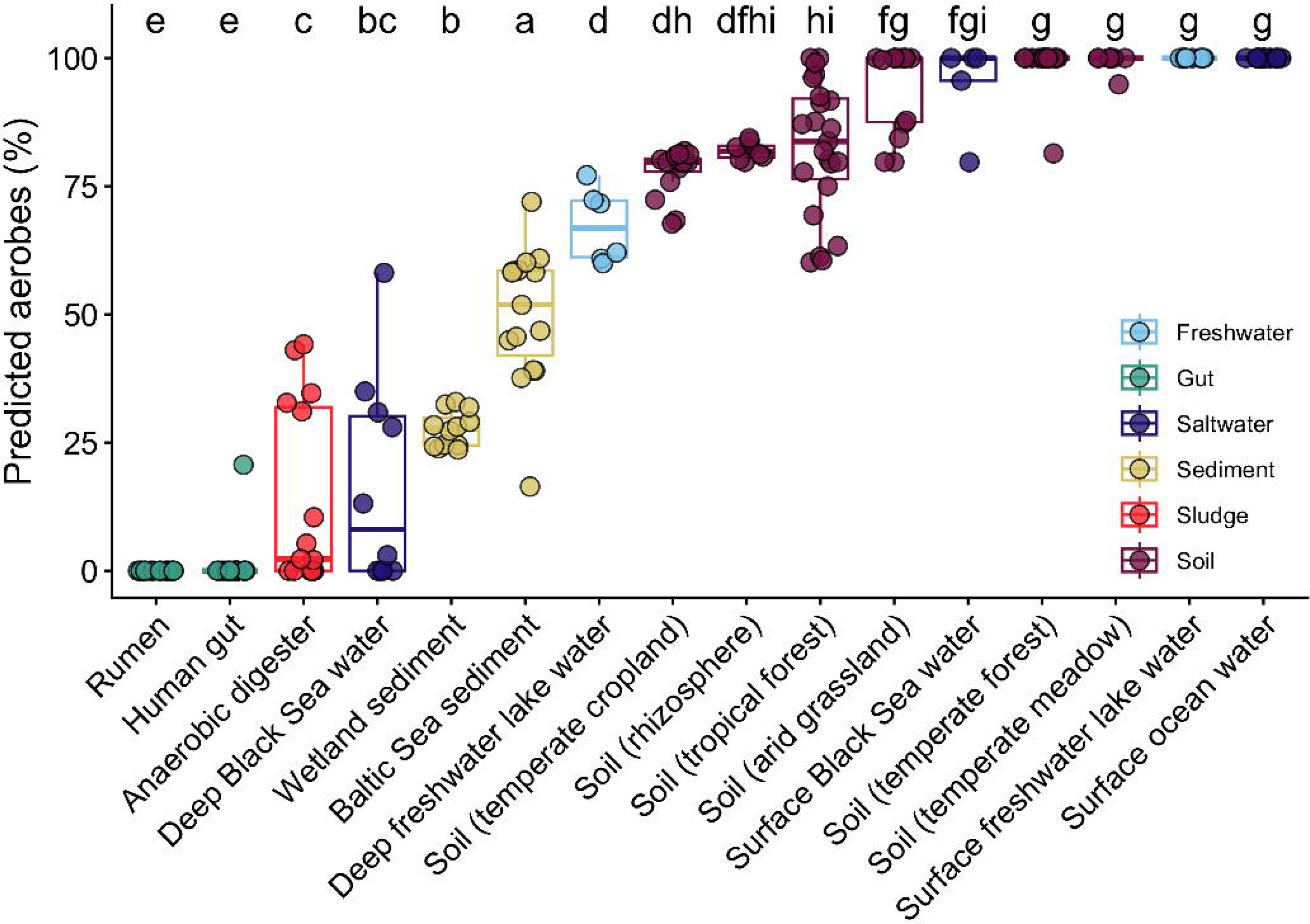
Predicted relative abundance of aerobic bacteria across 16 habitat types. The x-axis is sorted from left to right according to mean predicted percent abundance of aerobes. Different letters represent significant differences (Tukey posthoc, p < 0.05).

**Figure 4.**
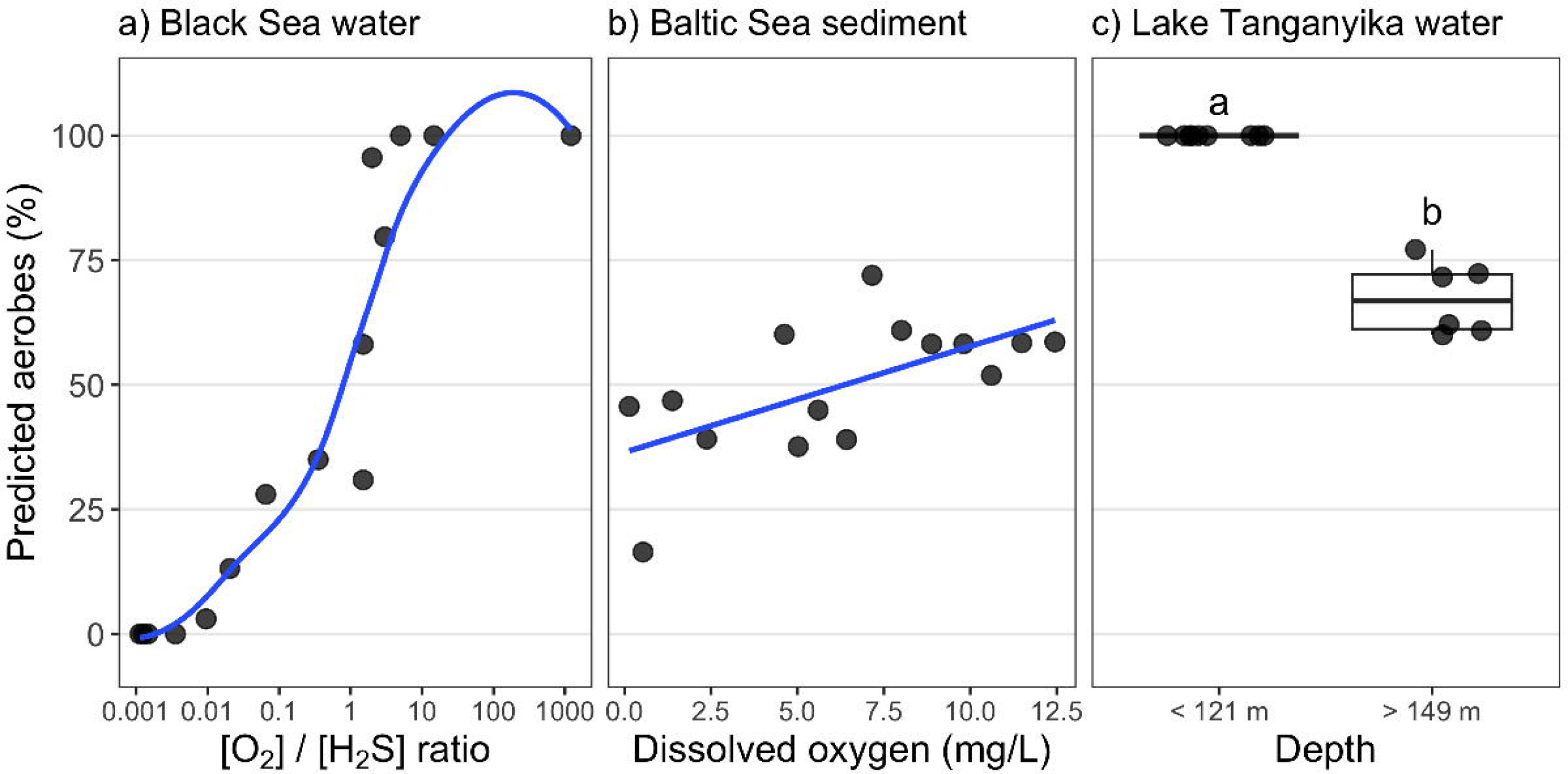
The predicted percent aerobe relative abundance significantly increased. a) with increasing ratio of oxygen to hydrogen sulfide across the Black Sea water depth gradient (GAM, p < 0.05), b) with increasing oxygen levels across sediments from the Baltic Sea (linear regression, p < 0.05), and c) in surface waters compared to deeper hypoxic waters in Lake Tanganyika (t-test, p < 0.05). Shown are a loess function in panel a, a linear model in panel b, and different letters representing a significant difference according to a t-test in panel c.

Next, we tested OxyMetaG across surface soils that are generally considered to be oxic but can have variable oxygen concentrations depending on moisture and texture, with some surface soils harboring anaerobes in microsites (21). Across 331 Australian soils, ranging from arid sites in western Australia to humid sites in Tasmania and northeastern Australia, the predicted relative abundance of aerobic bacteria ranged from 60% to 100% (mean = 95%, SD = 9%, median = 100%). The majority of these soils were dominated by aerobic bacteria, with 204 samples predicted to have 100% aerobes and 127 samples predicted to have some anaerobes.

Across 209 soils collected from 26 sites across the U.S., ranging from tropical to subarctic climates, the predicted relative abundance of aerobic bacteria ranged from 40% to 100% (mean = 95%, SD = 10%, median = 100%). Similar to Australian soils, most U.S. soils were dominated by aerobic bacteria, with 145 samples predicted to have 100% aerobes and 64 samples predicted to have some anaerobes. Ecosystem type significantly affected the proportion of aerobic bacteria in Australia (ZIBR, p < 0.001) but not in the U.S. (ZIBR, p = 0.22) (Figure 5). Climate class did not significantly affect the proportion of aerobes in Australia (ZIBR, p = 0.13) or the U.S. (ZIBR, p = 0.96) (Figure S3). As for continuous soil and climate variables, gravimetric water content was the only variable included in our models that was significantly associated with the predicted proportion of aerobes in both datasets (Table S3, Figure S4). Sand content, organic carbon content, and mean annual temperature were also significantly associated with the predicted proportion of aerobes in the Australian dataset (Table S3).

**Figure 5.**
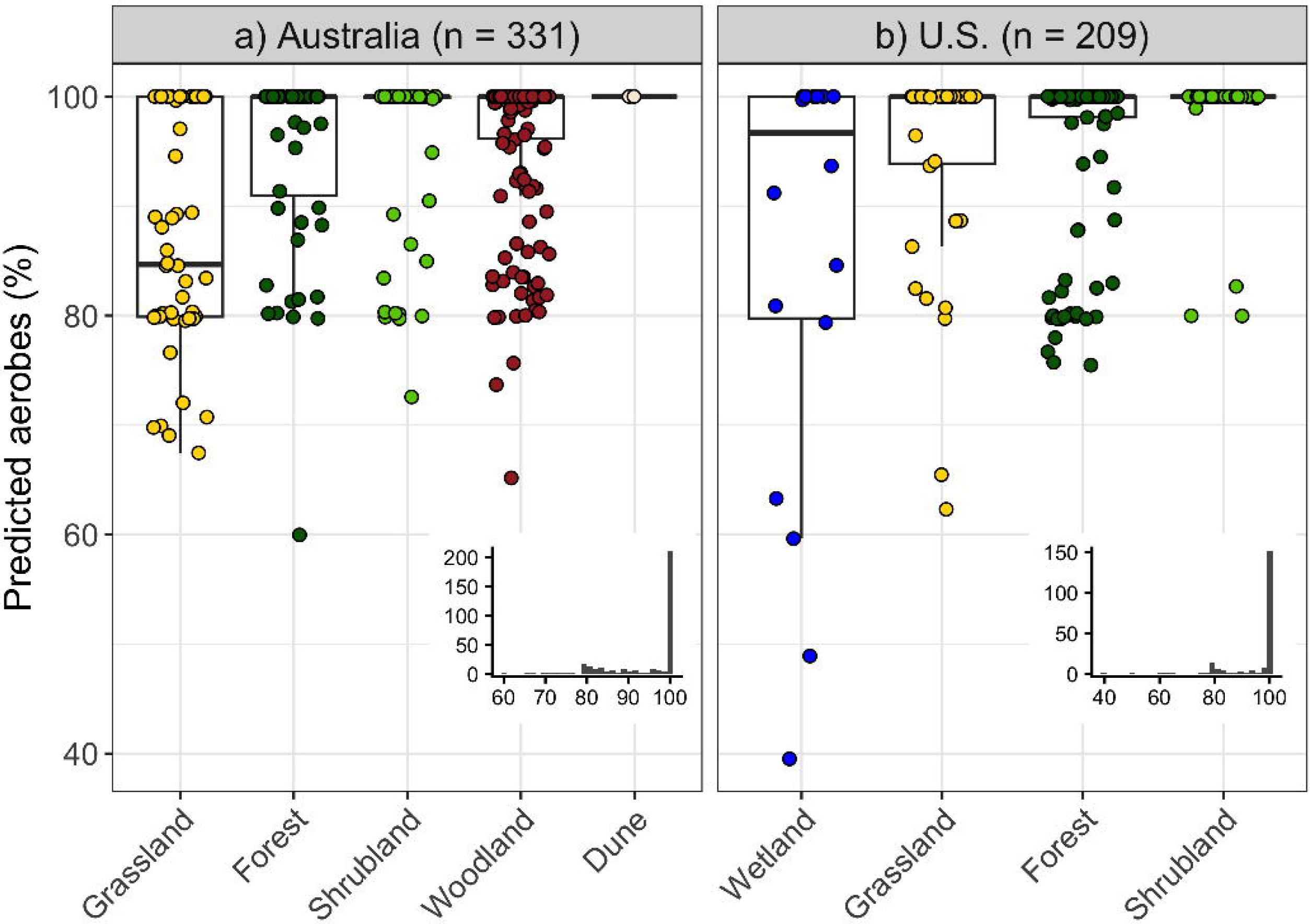
Predicted relative abundance of aerobic bacteria across habitats in. a) Australia and b) the United States. The x-axis within each panel is sorted from left to right by mean predicted percent aerobes. Insets show histograms; note the difference in axes scales between panels. Grassland, forest, and shrubland habitats were shared between the two datasets, while only Australia had dune samples and only the U.S. had wetland samples. Habitat significantly affected the predicted relative abundance of aerobic bacteria in Australia (ZIBR, p = 0.0002) but not in the U.S. (ZIBR, p = 0.22).

Lastly, we tested OxyMetaG with metagenomes from the human gut, which is generally considered to be anoxic, but can be oxic in the first few months of life (28). Across the 73 metagenomes tested, the predicted relative abundance of aerobic bacteria ranged from 0 to 61%. The proportion of aerobic bacteria declined significantly with time after birth (LMER, χ² = 18.9, p < 0.001), but there was considerable variation across individuals (Figure 6), with the model including random slopes and intercepts for subject IDs significantly outperforming the model with only random intercepts (ANOVA, χ² = 7.3, p = 0.03). All individuals eventually had 0% aerobic bacteria by the later time points, but the timepoint at which this occurred varied, with one individual still having aerobes past day 600 (Figure 6).

**Figure 6.**
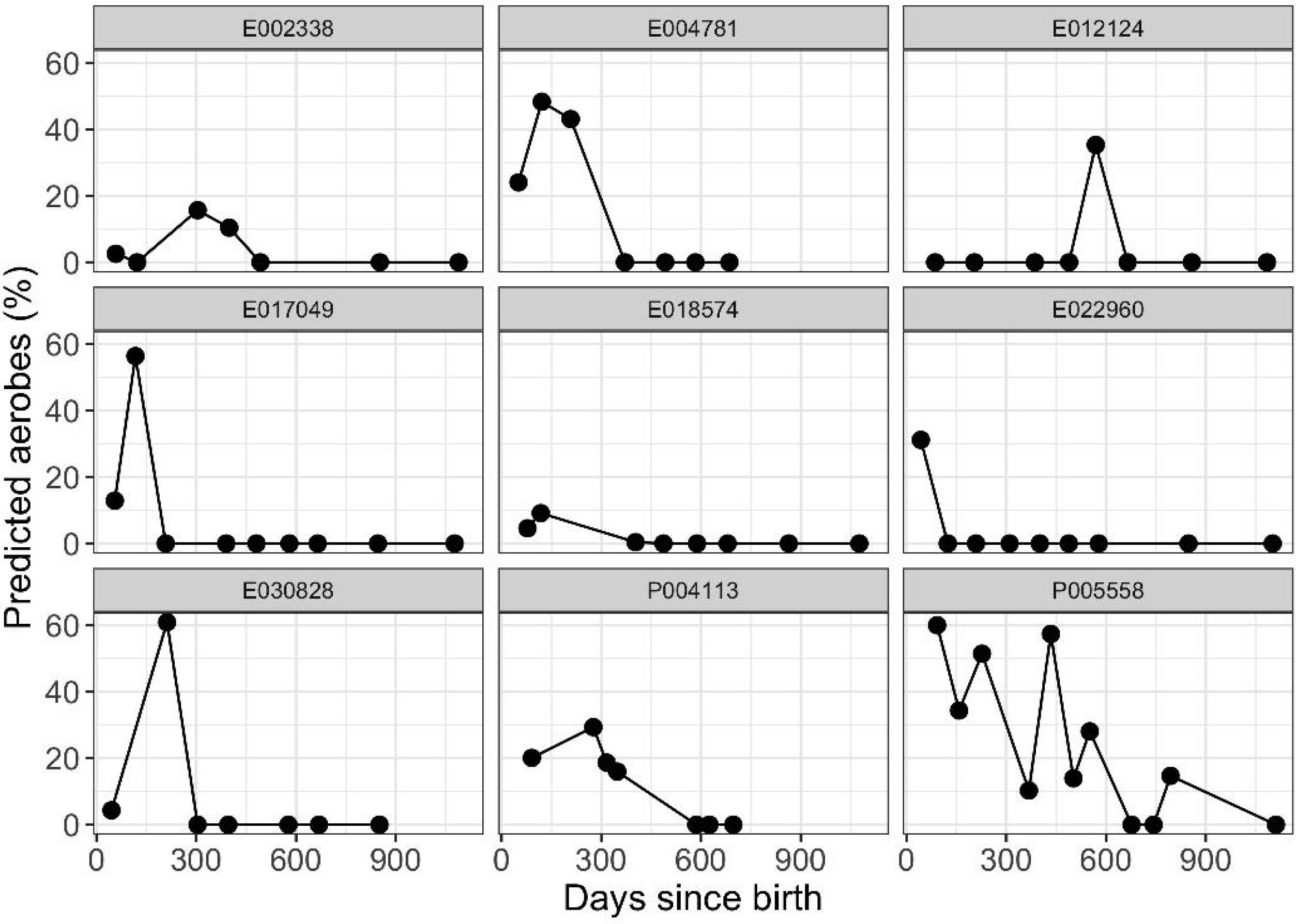
Predicted relative abundance of aerobic bacteria over time across 9 individuals from the DIABIMMUNE cohort study (66). The predicted relative abundance of aerobic bacteria significantly decreased over time (LMER, χ² = 18.9, p < 0.001).

## Discussion

We developed a metagenome-based tool, OxyMetaG, that uses a gene abundance ratio to infer the percent relative abundance of aerobic bacteria in a wide variety of environmental samples. OxyMetaG outputs the predicted percent relative abundance of aerobic bacteria (out of total bacteria) in a sample, which can be used as a bioindicator of general oxygen availability.

We identified 20 genes whose presence or absence could accurately predict whether a given bacterial taxon was aerobic or anaerobic across the majority of bacterial phyla (Figure S1, Table S1). These 20 genes are typically involved in various pathways of aerobic or anaerobic metabolic processes or in free radical scavenging, with a subset of these genes (9 of 20) having also been identified as predictors of bacterial oxygen tolerance in previous studies (6, 12–14). The gene abundance ratio was strongly related to the relative abundance of known aerobic bacteria in simulated metagenomes (Figure 2), demonstrating the predictive power of this metric. We first tested OxyMetaG on a diverse set of 203 terrestrial, aquatic, marine, and host-associated metagenomes. We then used OxyMetaG on 540 surface soils across two continents to investigate how the percent of aerobic bacteria varies across surface soil bacterial communities, and on 73 infant to toddler-age human gut samples to investigate patterns in the percent aerobic bacteria in the gut over time.

OxyMetaG accurately captured expected differences in community-level bacterial oxygen tolerances, predicting a dominance of anaerobes in known hypoxic to anoxic environments such as cattle rumens, human guts, anaerobic digesters, and wetland sediments, with aerobes found to dominate in environments that we typically consider to be oxic, such as surface soils and surface marine and freshwater samples (Figure 3). These results were similar to those obtained using other tools that rely on classifying individual genomes from those sample types (15, 16). One advantage of OxyMetaG compared to genome-based approaches is that it works on raw reads and does not require genome assembly, which can be a major limitation in many samples where obtaining metagenome-assembled genomes (MAGs) can be difficult, or when the MAGs that are obtained only represent a fraction of the diversity (17). Furthermore, in metagenomes from the Black Sea, the Baltic Sea, and Lake Tanganyika, the inferred changes in the relative abundances of aerobes within environments were well correlated with known oxygen gradients (29) (Figure 4). Together, these validation tests demonstrated how the OxyMetaG output (predicted percent aerobes) can be used for broad comparisons among sample types as well as more specific comparisons within sample types collected across gradients.

As a test case for demonstrating the utility of OxyMetaG for quantifying community-level patterns in bacterial oxygen tolerances across environmental gradients, we analyzed 540 metagenomes representing a broad range of soil types, habitat types, and climate conditions across Australia and the US. Most samples across both datasets were overwhelmingly dominated by aerobes, with OxyMetaG suggesting 100% relative abundance of aerobes in a majority (74% of samples). While these samples may in fact harbor a small proportion of anaerobic taxa, the high ratio (> 30) of aerobic indicator genes to anaerobic indicator genes in these samples suggests that any anaerobes may be relatively rare and temporally transient. These results suggest that, despite the presence of anaerobic microsites, surface soils generally select for, and are dominated by, aerobic bacteria. In line with predictions (21), soil moisture was the top variable associated with the percent aerobic bacteria in both datasets (Table S3), with wetter soils having slower diffusion of atmospheric oxygen (Figure S4). Still, the variation explained by soil moisture was low (Table S3), likely due to moisture being a single time-of-sampling measurement and the complexity of variables that influence soil oxygen availability.

As a second test case for OxyMetaG, we analyzed 73 human gut metagenomes, focusing on individuals that were sampled repeatedly over their first 3 years of life. Prior work using 16S rRNA marker gene sequencing combined with trait databases (matched by genus and species name or with hidden state prediction algorithms) showed that the abundance-weighted mean oxygen tolerance decreased markedly from months 3-15 (28). Results from our metagenome-based approach generally supports the idea of elevated oxygen availability in the gut during infancy (30), with a trend towards anaerobic dominance by year three that is likely driven by increases in *Bifidobacterium* and members of Bacillota such as Clostridiaceae (31, 32). Importantly, our data show a greater degree of variation than the prior work, likely due to the fact that OxyMetaG considers all bacterial reads and the gene-abundance-based method does not rely on trait databases biased towards well-characterized taxa.

One limitation of OxyMetaG is that we do not know the temporal scale that is integrated by our metagenome-based metrics. Standard metagenomic sequencing approaches do not discriminate between live and dead cells (33), nor do they capture variation in taxon-specific growth rates (34). Thus, OxyMetaG inferences of oxygen availability likely integrate over longer time periods that reflect both current conditions at the time of sampling and past conditions. This is beneficial in that OxyMetaG predictions are more representative of the sample history and general sample conditions, not just the single sampling moment, and the predictions are likely more robust to rapid variations in oxygen levels. The downside is that OxyMetaG would not be useful for studying changes in oxygen availability over short time scales. Although we currently do not know the exact time period that is integrated, as this will depend on community turnover rates, future work could investigate the temporal changes in aerobe versus anaerobe ratios in communities sampled over time where oxygen levels are experimentally manipulated. Another limitation of OxyMetaG is its omission of facultative aerobes/anaerobes and microaerophiles during training and gene identification; a useful next step would also be to incorporate genomic features from these taxa into OxyMetaG to more fully capture the spectrum of bacterial oxygen preferences.

We have presented results from OxyMetaG for a total of 816 modern metagenomic samples and expect OxyMetaG to be useful for inferring oxygen levels and answering ecological questions in more studies on modern metagenomes. Additionally, we also suggest that OxyMetaG could be useful in ancient metagenomic studies, where past oxygen levels are unknown and can only be inferred using geochemical proxies. OxyMetaG could be used in concert with other geochemical paleo-oxybarometers (35) for more robust inference about sample history. In a first pilot study on ancient metagenomes, OxyMetaG predictions were in line with isotopic data and helped identify two clusters of communities (aerobic and anaerobic) in subglacial lake samples from Antarctica (De Sanctis et al. 2025). OxyMetaG could contribute to the toolkit of paleo-oxygen proxies in that it is universal (as long as the sample has bacterial reads) and does not need to be calibrated on each individual sample or sample type. However, OxyMetaG still requires further tuning and optimization to handle issues specific to ancient DNA analyses, such as DNA damage and shorter fragment lengths; such improvements will be released in future updates to OxyMetaG. We hope OxyMetaG is ultimately useful and contributes to a better understanding of both microbial ecology and environmental conditions in general.

## Materials and Methods

Our workflow consisted of first modeling oxygen tolerance at the genome level to identify key genes that are highly predictive of oxygen tolerance (“indicator genes”), then using simulated metagenomes with known percentages of aerobic bacteria to establish a relationship between the percentage of aerobic bacteria and the ratio of aerobic indicator genes to anaerobic indicator genes, and then testing with actual metagenomes both for validation and application (Figure 1).

We first downloaded 5520 bacterial genomes from 31 different phyla that had oxygen tolerance information available in BacDive in June 2025 (36) (Table S4, Figure S5). Since our goal was to identify the most important genes associated with clearly aerobic and anaerobic bacteria, we only used taxa classified as “obligate aerobe”, “aerobe”, “anaerobe” or “obligate anaerobe”, and omitted taxa classified as “facultative aerobe”, “facultative anaerobe”, or “microaerophile”. We acquired additional information about the genomes, including taxonomic information, by matching accessions to GTDB r226 (37). We used hmmer v3.4 (38) to annotate the genomes with the Pfam v37.3 database (39). We then used an ensemble machine learning approach to select the top 20 genes most predictive of oxygen tolerance. This approach uses CatBoost, XGboost, and RandomForests, an 80:20 ratio of training to testing data, and SHAP scores to assess variable importance (40). After building the best model using all genes, the model was rerun with just the top 20 genes with the highest SHAP scores. The model with only 20 genes performed as well as the model with all genes in terms of precision, recall, and F1-score (weighted averages all 0.96 for both models), identifying 13 genes associated with aerobes and 7 with anaerobes (Table 1, Figure S1). The prevalence of the 13 aerobic indicator genes ranged from 54% to 99% in aerobic taxa and from 2% to 47% in anaerobic taxa, while the prevalence of the 7 anaerobic indicator genes ranged from 51% to 89% in anaerobic taxa and 1% to 17% in aerobic taxa (Table 1). Furthermore, the accuracy of the model was high even when tested on individual phyla (F1 > 0.90). Tests were conducted on each of the four phyla with the most genomes (Pseudomonadota, Actinomycetota, Bacillota, Bacteroidota), as well as on all of the remaining 27 phyla combined. Furthermore, accuracy of the main model (trained using all phyla, tested using all phyla) was assessed for each phylum to quantify model performance per phylum. The scripts to implement our approach are publicly available on Zenodo (https://doi.org/10.5281/zenodo.18331946).

We then used ART (41) to simulate metagenomes (Illumina, 150 bp) using 500 randomly selected genomes from the original set of 5520 genomes. To test for any effect of sequencing depth, simulations were done using fold coverages of 0.5 and 2.5, which generated 5-8 million reads and 26-41 million reads, respectively, depending on the randomly selected subset of genomes used in the simulation. We used a gradient of 0, 20, 40, 60, 80, and 100% of the genomes as aerobes. We used three replicates per aerobe percentage, with the specified number of aerobic and anaerobic genomes randomly selected from the available pool of 5520 genomes. Reads were trimmed and quality filtered with Trimmomatic v0.39 to remove Illumina adapters and retain reads with Q ≥ 20 and length ≥ 100 bp (42). We then downloaded all of the reviewed protein sequences for each Pfam as a fasta file from the Pfam website, concatenated them into a single fasta file, and generated a DIAMOND database with DIAMOND v2.1.13 (43). We then used DIAMOND BLASTX to calculate the abundance of the 20 genes in the metagenomes. We retained hits with percent identity ≥ 60, e-value < 0.001, and bitscore ≥ 50 (44). We then normalized by mean Pfam length to calculate reads per kilobase (RPK) for each Pfam. We summed the RPK of the 13 aerobic genes and the 7 anaerobic genes and then calculated the aerobic: anaerobic gene RPK ratio. Because the relationship was nonlinear, we fit a generalized additive model (GAM) of the relationship between the percent aerobe genomes and the ratio (Figure 2) using the mgcv R package (45). The model relationship was highly similar across the two sequencing depths tested (Figure 2).

We then validated the model on 203 modern metagenomes that we expected to capture broad differences in oxygen concentrations, including a subset of samples with paired measurements of oxygen concentrations. The 203 metagenomes spanned 16 distinct environments, including cattle rumen (46), human gut (47), surface ocean water (top 5 m) (48), Black Sea water (surface and deep) (49), Baltic sea sediment (0-2 cm below sea floor) (50), anaerobic sludge digester (publicly available but unpublished), wetland sediment (51), freshwater lake water (surface and > 150 m deep) (29), surface soil (0-10 cm) from arid grasslands in Australia (52), surface soil (0-10 cm) from tropical forests in Panama (53), other soils ranging from 0 to 70 cm depth from temperate forests, grasslands, and meadows (54), and surface bulk and rhizosphere soil from maize crops (55). Approximately 15 samples were selected from each study (Table S5). For further validation, we conducted additional analyses on a subset of samples with measured gradients in oxygen concentrations. We compared 15 metagenomes from the Black Sea that spanned a depth profile from 50 m to 2000 m, with measured oxygen and hydrogen sulfide concentrations (49), 15 metagenomes from Baltic Sea sediments that spanned dissolved oxygen levels from 0.13 to 12.45 mg L^-1^ (50), and 15 freshwater lake metagenomes ranging from 0 m to 1200 m depth with oxygen saturation levels ranging from 1.3 to 91.4% (29). We downloaded raw metagenomic data from the NCBI SRA and trimmed and filtered the reads with Trimmomatic as above. Because not all of the 20 predictor Pfams are exclusive to bacteria, we extracted only bacterial reads from the metagenomes, as hits to archaeal or eukaryotic reads could introduce errors in our model. We used Kraken2 v2.1.3 (56) to classify all of the quality-filtered metagenomic reads and then KrakenTools (57) to extract only the bacterial reads from each metagenome. The number of bacterial reads per metagenome averaged 10.9 million, with only four samples below 256000 reads. Then, as above, we used DIAMOND BLASTX to calculate the abundances of each Pfam and then calculated the RPK ratio of aerobic indicator genes to anaerobic indicator genes to predict the relative abundance of aerobic bacteria in the sample. Predictions over 100% were set to 100%, predictions below 0% were set to 0%, and any ratios above 30 (the maximum value in simulations) were set to 100%.

OxyMetaG is implemented as a Python package with three main functions, one for each of the steps outlined above. The ‘extract’ function is a wrapper for Kraken2 and KrakenTools, which together classify each read and then extract reads classified as bacterial. The ‘profile’ function takes the bacterial reads as input, runs DIAMOND BLASTX on the database of the 20 Pfams, and records the information about hits for each sample. Lastly, the ‘predict’ function takes the DIAMOND output files, filters them to the top significant hit per read, calculates the summed aerobic: anaerobic RPK ratio, and predicts the proportion of aerobic bacteria using the GAM established with the simulated metagenomes. While we used cutoffs of percent identity ≥ 60, e-value < 0.001, and bitscore ≥ 50, users can also specify their own percent identity, e-value, and bitscore cutoffs based on their specific data characteristics, such as read length, and desired balance in sensitivity and specificity (58). This mode also enables users to test for consistency of results across multiple tested cutoff values.

To test the effect of sequencing depth on the predictions, we chose one oxic sample, one anoxic sample, and one mixed sample that each had high sequencing depth (>32 million reads per sample). We randomly subsampled each of the three metagenomes to 16 different sequencing depths (ranging from 0.001 to 32.768 million reads) running OxyMetaG separately at each sequencing depth. The predicted relative abundance of aerobic bacteria stabilized at only 256000 reads, demonstrating the utility of this method for analyzing metagenomes with relatively low sequencing depths that would preclude assembling genomes (Figure S6). This was also the point at which there were anaerobic genes detected in the oxic sample, enabling the ratio to be calculated and the prediction to be made (Figure S6).

After validation, we applied OxyMetaG on soil and human gut datasets to answer ecological questions in those systems. First, we used 331 surface soil (0-10 cm) metagenomes collected using standardized methodology from across Australia (52). These metagenomes have been previously used in other research (59) and were selected because they are from natural areas, had at least 10 million reads per metagenome, and had defined upland vegetation types. Quality-filtered shotgun metagenomes (150 bp) and metadata (Table S6) were downloaded from the Bioplatforms Australia data portal (https://data.bioplatforms.com/bpa/otu/metagenome).

Mean annual precipitation and mean annual temperature were determined for each sample using the WorldClim2 database (60), and aridity index was determined using the Global Aridity Database version 3 map at 30 arc-second resolution (61). The number of bacterial reads in these Australian metagenomes ranged from 3 to 19 million (mean = 7.9 million). For comparison and to include more humid sites, we used an additional, independent dataset of 209 surface soil (0-30 cm) shotgun metagenomes (Table S7) collected by the National Ecological Observatory Network (NEON) using standardized methodology from 26 sites across the United States. We used samples collected in 2023 and sequenced at the U.S. Department of Energy Joint Genome Institute (JGI). Because OxyMetaG is assembly-free and does not require high sequencing depth (Figure S6), we subsampled all metagenomes down to 5 million reads before running OxyMetaG to reduce computational costs. The number of bacterial reads ranged from ∼1.1 to 2.2 million reads (mean = 1.7 million). Climate, soil, and other general metadata associated with each metagenome were downloaded using a combination of the phyloNEON R package (62) and neonUtilities R package (63). For statistical analyses, we converted the percent aerobe values, which had a highly skewed distribution towards 100% values, to a proportion anaerobe value, for use as a response in zero-inflated beta regression (ZIBR), where the response must be bounded between 0 and 1 (64). Relationships between the predicted proportion of anaerobes and ecosystem type, climate class, and continuous climate (mean annual precipitation and temperature, aridity index) and soil variables (gravimetric water content, pH, total carbon, total nitrogen, carbon: nitrogen ratio, nitrate, phosphorus, percent sand) were tested with ZIBR implemented in the gamlss R package (65). Models including each predictor variable were compared to a null model with the likelihood ratio test.

Second, we used 73 human gut metagenomes from Russia and Finland collected with standardized methodology for the DIABIMMUNE cohort study (66). From all of the available metagenomes in the study, we selected metagenomes from 9 individuals that were sampled across at least 7 time points, had the first sample before the 93rd day of life (minimum days after birth was 43), and had a sampling time range of at least 634 days (maximum day after birth was 1111) (Table S8). These metagenomes had an average of 1.5 million bacterial reads. We ran linear mixed effect regression (LMER) models in the lme4 R package (67) between predicted relative abundance of aerobic bacteria and days after birth. We compared two models with the ‘anova’ function in R: one that only allowed random intercepts for each subject ID, and one that allowed random intercepts and slopes for each subject ID. This enabled us to assess the effect of time (days since birth) and subject ID on the predicted relative abundance of aerobic bacteria. All downstream statistical and graphical analyses were performed in R 4.5.1 (68).

## Acknowledgements

We thank the NovoNordisk Foundation and Wellcome Trust for funding. We thank Hugh Cross for assistance in acquiring the NEON metagenomes and metadata. We thank Antonio Fernandez-Guerra for feedback on OxyMetaG.

## Data availability

The 5520 genomes used in the genome-level models are publicly available on NCBI, with accessions and metadata listed in Table S4. The 203 shotgun metagenomes used for the cross-habitat validation are publicly available on NCBI, with accessions and metadata listed in Table S5. The 331 BASE metagenomes from Australia are publicly available on the Bioplatforms Australia data portal (https://data.bioplatforms.com/bpa/otu/metagenome); sample IDs and metadata are provided in Table S6. The 209 NEON metagenomes from the United States in 2023 are publicly available on the JGI data portal (https://data.jgi.doe.gov/) under JGI_ID 509938, with “2023” contained in the sample name; sample IDs and metadata are provided in Table S7. The 73 human gut metagenomes are a subset of the publicly available DIABIMMUNE metagenomes available on NCBI BioProject PRJNA290380; exact accessions and metadata are detailed in Table S8.

## Code availability

OxyMetaG is available from GitHub (https://github.com/cliffbueno/oxymetag) and PyPI. Data analysis scripts for this manuscript can be found on Zenodo (https://doi.org/10.5281/zenodo.18331946).

