## Supplementary material for "Quantifying the oxygen preferences of bacterial communities using a metagenome-based approach": Figure S

Running title: Quantifying bacterial oxygen preferences

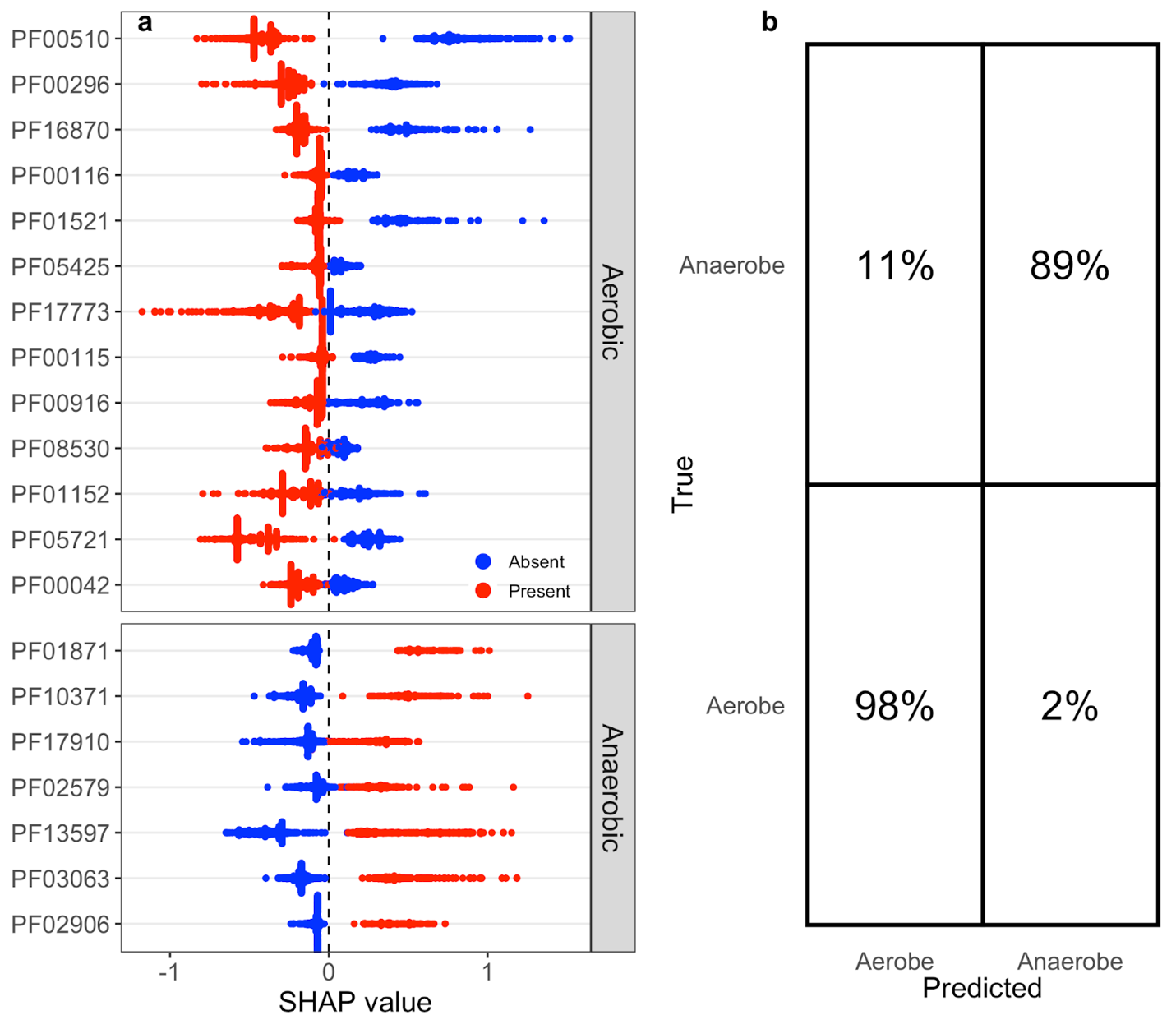

Figure S1. a) Average SHAP values of the 20 selected Pfams (13 aerobic indicators, 7 anaerobic indicators) from ensemble machine learning classification modeling. For Pfam names and other information, see Table 1. b) Confusion matrix of the genome-level ensemble machine learning model predicting whether a genome comes from an aerobe or anaerobe based on BacDive data (training n = 4416, testing n = 1104).

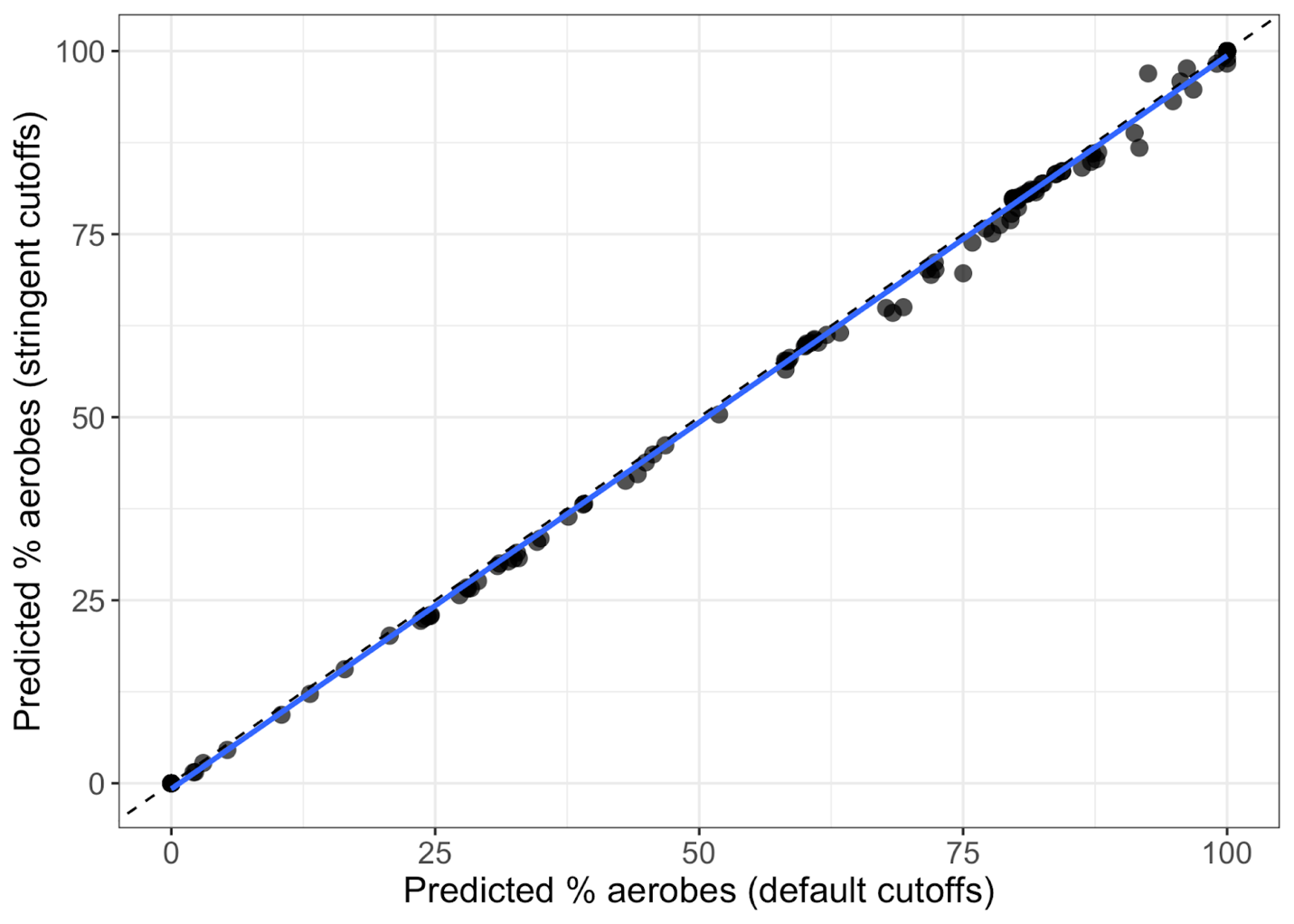

Figure S2. Comparison of the predicted relative abundance of aerobic bacteria across the 16-habitat dataset (Figure 4) using two different DIAMOND BLASTX hit cutoffs. The default cutoffs . n = 173 of the 203 samples from Figure 4 that had predictions using both cutoffs (with stringent cutoffs, there were 20 samples that did not have at least one aerobic indicator Pfam and at least one anaerobic indicator Pfam). The dashed black line represents the 1:1 line and the solid blue line is the linear regression line. The two predictions were strongly and significantly positively correlated (R^2^ = 0.99, p < 0.001). Default cutoffs: percent identity ≥ 60, e-value < 0.001, and bitscore ≥ 50. Stringent cutoffs: percent identity ≥ 60, e-value < 1e-10, bitscore ≥ 60, and amino acid alignment length ≥ 40.

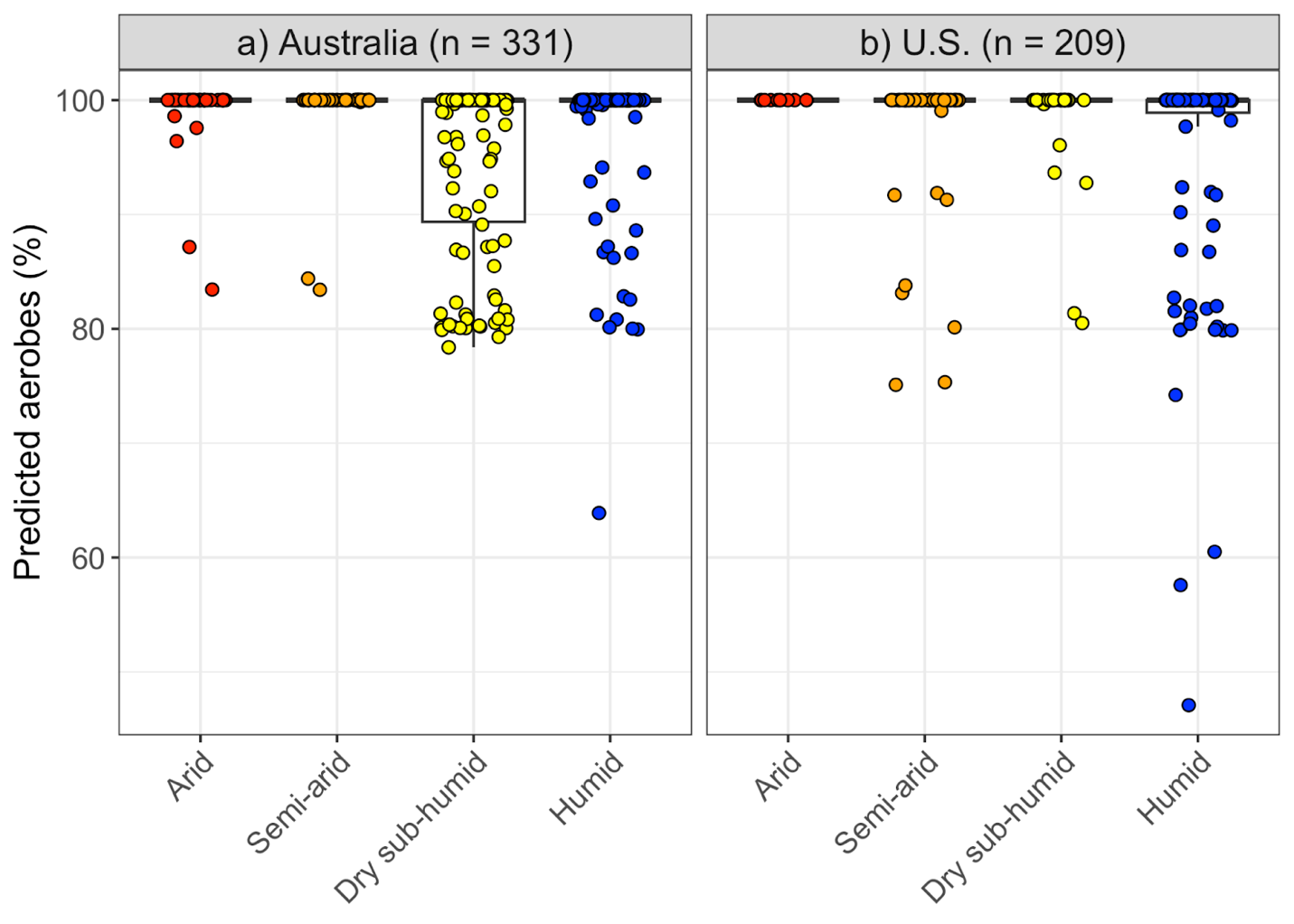

Figure S3. Predicted relative abundance of aerobic bacteria across climate classes in a) Australia and b) the United States. The x-axis within each panel is sorted from left to right by mean predicted percent aerobes. Insets show histograms; note the difference in axes scales between panels. Climate class did not significantly affect the predicted relative abundance of aerobic bacteria (ZIBR, p > 0.05).

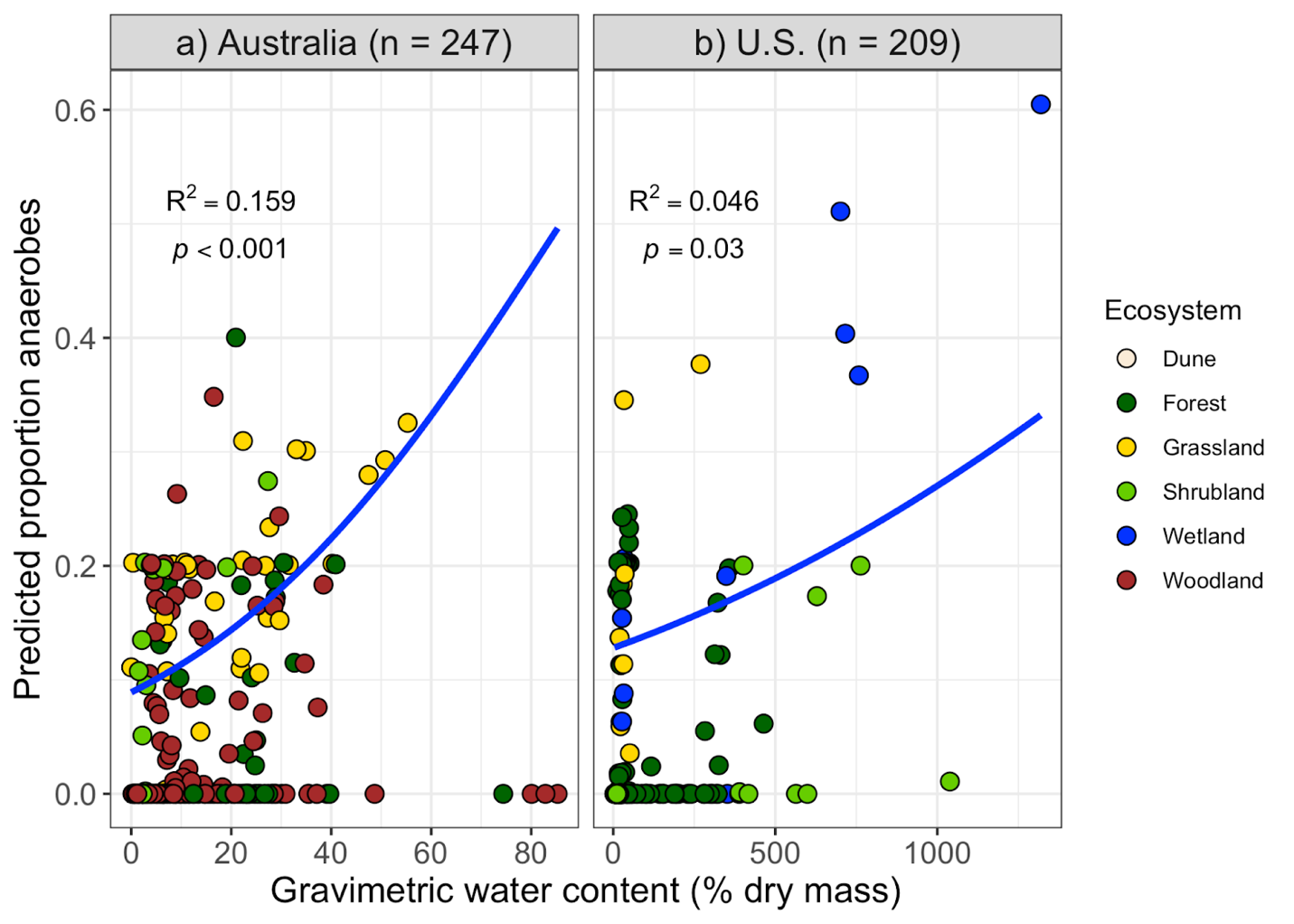

Figure S4. The predicted proportion of anaerobes was significantly positively associated with gravimetric water content in both datasets according to zero-inflated beta-regression (p < 0.05). Note that of the 331 Australian soils analyzed in this study, only 247 included data on gravimetric water content and are included in this figure.

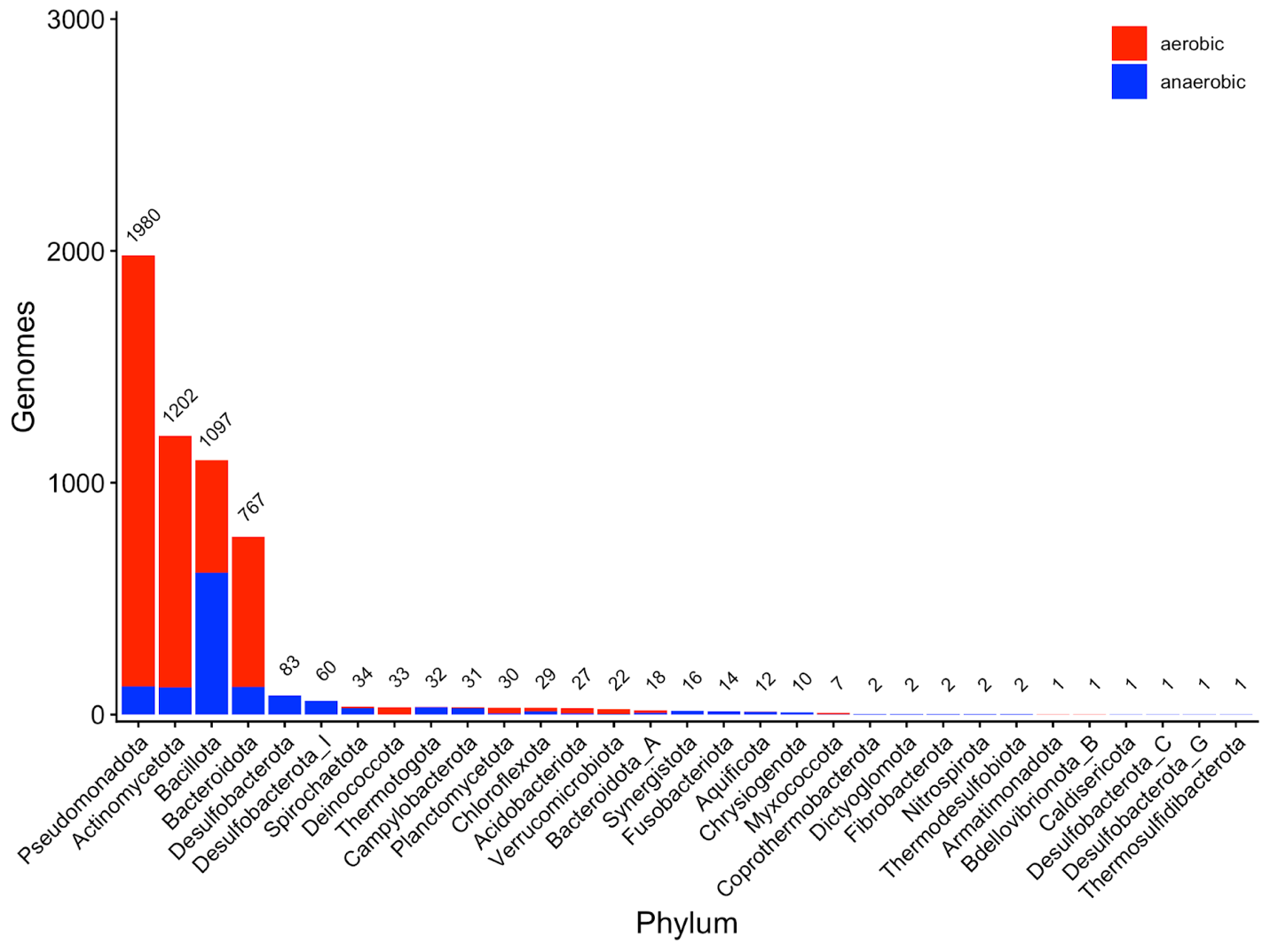

Figure S5. Number of genomes of each phylum (according to GTDB taxonomy) and each oxygen tolerance category (BacDive), n = 5520. These genomes were inputs in an ensemble machine learning model to predict oxygen tolerance from Pfam presence. Numbers above the bars represent the number of genomes in each phylum.

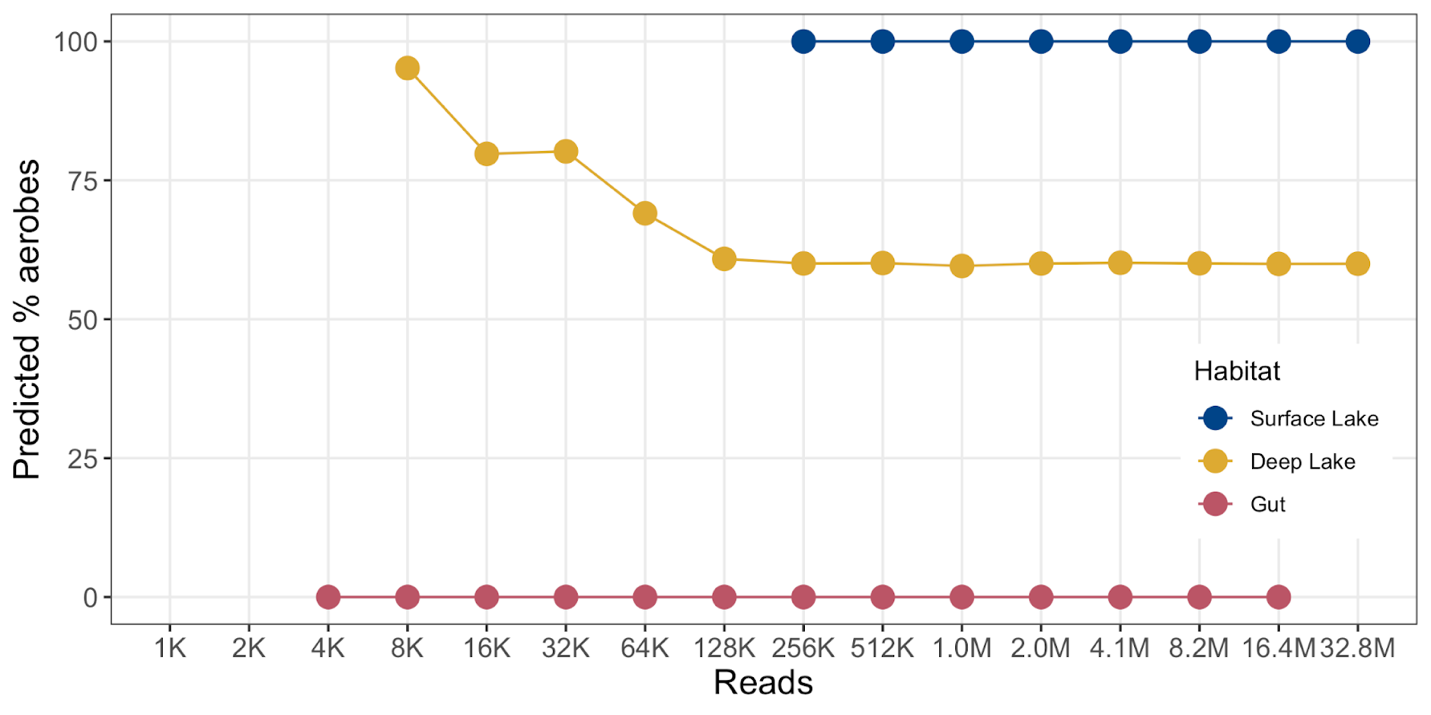

Figure S6. Assessment of the effect of sequencing depth on the predicted relative abundance of aerobic bacteria. No prediction is made if 0 aerobic Pfams or 0 anaerobic Pfams are detected, which was the case until 4k reads for the human gut sample, 8k reads for the deep lake sample, and 256k reads for the surface lake sample. Note that the human gut sample did not have 32.8 million bacterial reads.
